# Functional Subtypes of Synaptic Dynamics in Mouse and Human

**DOI:** 10.1101/2023.05.23.541971

**Authors:** John Beninger, Julian Rossbroich, Katalin Tóth, Richard Naud

## Abstract

Synapses show preferential responses to particular temporal patterns of activity. Across individual synapses, there is a large degree of response heterogeneity that is informally or tacitly separated into classes, and typically only two: facilitating and depressing short-term plasticity. Here we combined a kernel-based model and machine learning techniques to infer the number and the characteristics of functionally distinct subtypes of short-term synaptic dynamics in a large dataset of glutamatergic cortical connections. To this end, we took two independent approaches. First, we used unsupervised techniques to group similar synapses into clusters. Second, we used supervised prediction of cell subclasses to reveal features of synaptic dynamics that characterized cellular genetic profiles. In rodent data, we found five clusters with a remarkable degree of convergence with the transgenic-associated subtypes. Two of these clusters corresponded to different degrees of facilitation, two corresponded to depression with different degrees of variability and one corresponded to depression-then-facilitation. Strikingly, the application of the same clustering method in human data inferred highly similar clusters to those observed in rodents, supportive of a stable clustering procedure and suggesting a homology of functional subtypes across species. This nuanced dictionary of functional subtypes shapes the heterogeneity of cortical synaptic dynamics and provides a lens into the basic motifs of information transmission in the brain.

## Introduction

Studying how neurons communicate information provides a window into the fundamental language of the brain. This language is partly articulated by synapses that show preferential responses to particular temporal patterns of activity. This selectivity is enabled by transient (10 ms - 10 s) changes in connection strength termed short term plasticity (STP) (1–3). STP is found in most brain regions and helps sculpt neural information processing in-vivo (4, 5). The most basic types of STP are short term facilitation (STF) and short term depression (STD) which entail a transient increase or decrease in synaptic strength, respectively (6–9). More complex forms of STP have been reported, including biphasic STP (10, 11), delayed burst-dependent STP (12), and sub- and supra-linear facilitation (13). The prevalence and complexity of STP highlights the importance of understanding its role in shaping information transmission (14).

Short term plasticity has long been viewed as a mechanism to allow for communication of different messages to different postsynaptic targets from the same axon (15–18). Depending on the timescale of STP, it supports detection of subtle changes in low versus high firing rates (19–21) or subtle changes in firing patterns such as bursts versus single spikes (20). A number of other network-level functions have also been advanced: Depressing synapses may serve to decorrelate inputs (22) while facilitating synapses can enhance detection of correlated signals (23). STD can be seen as an adaptation mechanism (24), allowing for preservation of time-delay information across multiple levels of intensity (14, 25, 26). STP has also been proposed to support working memory either by serving directly as its basis (27) or by acting at a network level to sculpt the emergence of persistent attractor dynamics (28). In addition, STP may facilitate network-level storage and retrieval of memories (29). In spike trains encoding a multiplexed code, STP properties can be used to decode independent streams of information (30–32). Decoding multiplexed code is broadly relevant to biological learning because this type of multiplexed coding has been used to address the “credit assignment problem”, providing an account of how learning can be coordinated across multiple areas (32–35). Finally, the diversity of STP across individual synapses can form a reservoir of temporal patterns, which is beneficial to solving temporal credit assignment problems that involve learning temporal associations (36, 37). This myriad of roles of STP underscore the importance of clearly understanding the functional subtypes of STP that exist in biology.

A major impediment to relating STP to its function is the large degree of response heterogeneity that exists across synapses (38, 39). This heterogeneity is routinely and tacitly separated into broad classes such as facilitation and depression. A number of studies have established that, even when controlling for post-synaptic neuron subclass, this heterogeneity appears to be structured (37–40). While this heterogeneity arises from a combination of intrinsic variability, plasticity and genetically regulated synapse types, how these factors come together remains a matter of debate. Three hypotheses can be proposed: 1) STP properties walk a continuum, with cell types associated to parts of the continuum, 2) STP properties are clustered by genetically defined functional subtypes, with intrinsic variability and plasticity acting in a small neighborhood around the cluster center, 3) STP properties cluster around genetically defined subtypes, but a continuum of STP properties between subtypes are allowed. A prediction of hypotheses 2 and 3 but not of hypothesis 1 is that functional subtypes should emerge from an unsupervised clustering of the dynamics of a large number of synaptic responses. A strong test of the validity of such clusters is their stability across species. Furthermore, determining the predictive power of genetic information on the synaptic dynamics can determine the portion of the variance that is genetically defined (11, 16–18, 38, 41). Theoretically, clusters found by unsupervised methods should also contain features that are predictive of the genetic makeup of cells on both sides of the synapse. These questions can be tested in a sufficiently large dataset and are the main focus of this article.

Here we leverage the Allen Institute Synaptic Physiology Dataset (42), which includes an unprecedented number paired-patch recordings performed under physiological calcium conditions. Crucially, this dataset provides enough synapses to meet the training requirements of data efficient machine learning techniques and open a new avenue of analysis. We have applied a highly adaptable short-term plasticity model (referred to here as the Spike Response Plasticity (“SRP”) model) (43) to generate empirical fits descriptive of STP. Using unsupervised learning across the 224 excitatory connections, we found 5-6 stable clusters of STP with a striking match between human and mouse. We then used the cell subclass labels provided by the Allen institute for supervised analysis of STP properties and contrasted the features used for supervised (label aware) vs unsupervised (label free) learning. Our findings suggest that a minimum of five functional STP subtypes form the biologically plausible building blocks of the information processing properties of synapses, and that these STP subtypes only weakly relate to the cellular subclass level of transcriptomic description.

## Results

Our analysis focused on glutamatergic cell connections in mouse and human cortex from the Allen Institute Synaptic Physiology Dataset (42). In this publicly available dataset, paired-patch recordings were performed by stimulating the pre-synaptic neurons with diverse temporal patterns and recording the excitatory post-synaptic amplitude. The experiments were performed with genetically-identified cells for both pre- and post-synaptic partners via transgenic mouse lines. The choice of genetic markers was chosen according to the subclass level of hierarchical clustering of transcriptomic data (44–46).

### Model captures the diversity of synaptic dynamics

In order to generate a biologically plausible dictionary of functional STP models, we fit instances of the Spike Response Plasticity (SRP) model from Rossbroich et al. (43) to synaptic data from the Allen Institute Synaptic Physiology Dataset (47). As shown in Figure 1A, the SRP model predicts post-synaptic efficacy from an input spike train while incorporating short-term changes in synaptic strength. The SRP model works by convolving an input spike-train with a spike-response kernel, taking a nonlinear readout of the resulting trace at the time of the spikes, and then stochastically sampling synaptic efficacies according to this readout. The spike-response kernel encapsulates the trace left by every spike on the degree of facilitation/depression. It can combine facilitation and depression on different timescales (Fig. 1A, right). The parameters of this model regulate the shape and baseline of the spike-response kernel as well as the standard deviation of the stochastic sampling step.

**Fig. 1.**
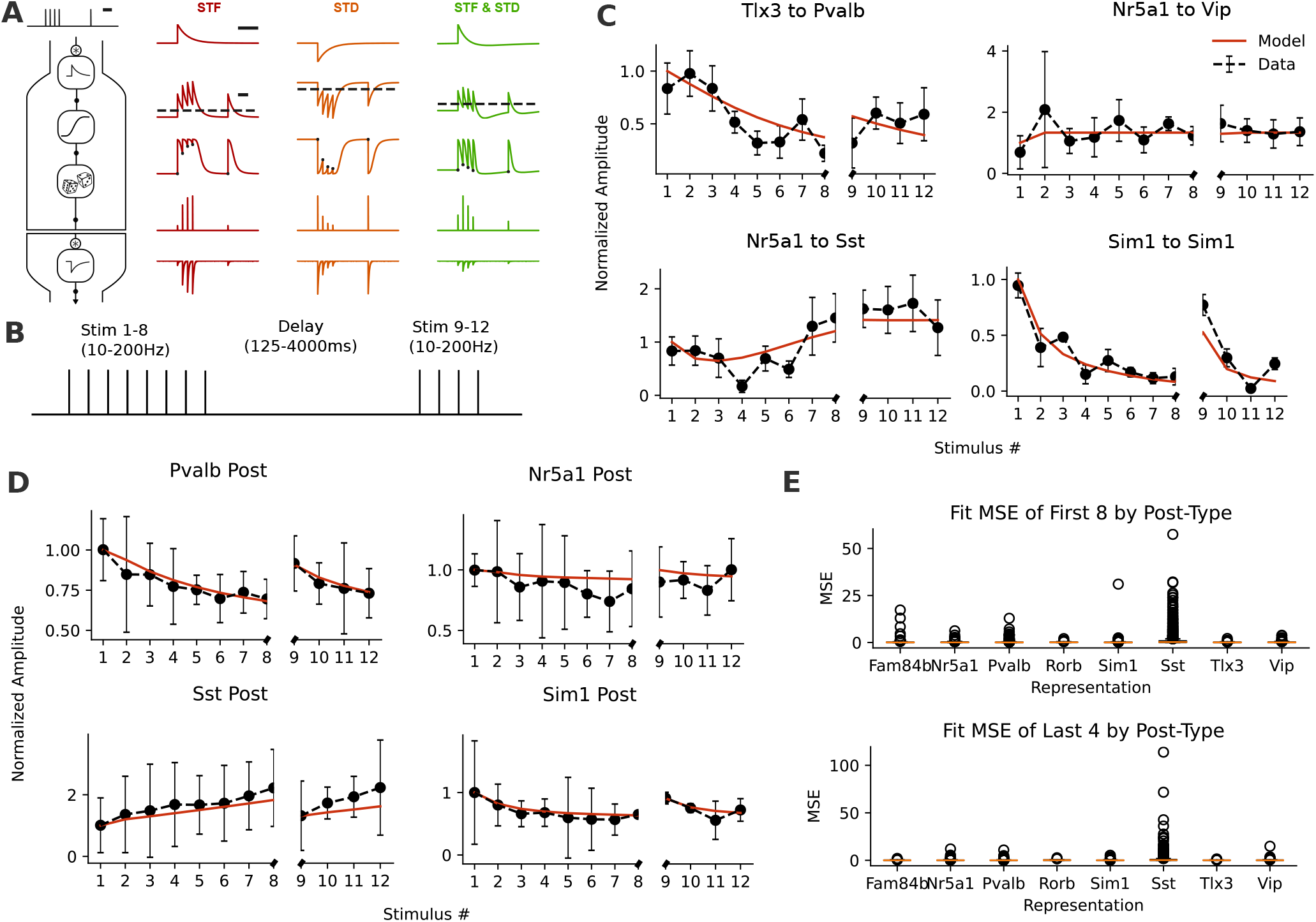
The Spike Response Plasticity (SRP) model captures responses across a diversity of synapse types. **A**: shows the SRP model of short-term synaptic dynamics from Rossbroich et al. (2021) (43) predicts excitatory post-synaptic potentials (EPSPs) based on spike timing input. **B**: shows the timing of input given to pre-synaptic cells in a paired patch clamp experiment, the model in A is fitted to predict EPSPs from this input. **C**: Shows fits of example synapses between different transcriptomic-associated subclasses for a particular protocol. The Tlx3 to Pvalb example corresponds to 20 Hz stimulation frequency and a 250 ms delay, the Nr5a1 to Vip example is for 50 Hz stimulation frequency and a 1003 ms delay, the Nr5a1 to Sst example is for 50 Hz stimulation frequency and a 125 ms delay, and the Sim1 to Sim1 example is for 50 Hz stimulation frequency and a 250 ms delay. **D**: shows mean SRP model fits (red), and data (black) for all synapses onto specific post-synaptic cell types (29 onto Pvalb, 11 onto Nr5a1, 11 onto Sst, 25 onto Sim1) and demonstrates high overall fit quality. Error bars show standard error of the mean (SEM). **E**: shows mean squared error of fit to individual runs for the first eight stimuli before a delay (top) and the four stimuli after a delay (bottom), grouped by postsynaptic cell subclass. The orange line indicates the median mean squared error while circles show outliers.

In order to capture the differing dynamics present within the Allen Institute Synaptic Physiology Dataset, we used standard optimization methods to fit distinct kernel parameters for every cell-pair. Figure 1B illustrates the range of stimulus protocols for which data were available. In all cases, eight stimuli were applied pre-synaptically at a fixed frequency, followed by a delay and four more fixed frequency stimuli. A range of stimulation frequencies and delay durations were experimentally examined for each synapse. Model parameters were optimized via maximum likelihood on the sequence of recorded post-synaptic amplitudes. The model captured both facilitating and depressing synapses, as shown in Fig. 1C for four exemplar pairs. We also found strong average fitting performance when we pooled protocols and pairs that corresponded to the same post-synaptic cell subclass (Fig. 1D). For each post-synaptic subclass, the mean squared error remained low (mean MSE and standard error of the mean of 0.38 ± 0.02 and 0.47 ± 0.06 for the first eight stimuli and the last four stimuli across all post-subclasses, respectively; Fig. 1E). Synapses onto somatostatin (Sst) expressing cells were notable for having a higher proportion of poorly fitting outliers. Specifically, the mean MSE and standard error of the means of all synapses onto somatostatin expressing cells was 1.4 ± 0.1 (p<0.001) for the eight pre-delay simulations and 1.9 ± 0.4 (p<0.001) for the four post-delay simulations. As no particular subclass apart from Sst was associated with worse average fits, the model parameters can be used as a compressed representation of the dynamics (48).

### Unsupervised clustering reveals functional subtypes

In order to develop new insights about the functional subtypes of STP in rodents, we used unsupervised machine learning techniques to perform clustering (2A). Clustering allows us to group synaptic dynamics based solely on the relative distance of synapses in a space. Instead of directly using the parameter space, we further improve the separability of the representation by focusing on the corresponding principal components. We selected OPTICS (Ordering Points To Identify the Clustering Structure) (49) as a clustering algorithm, which defines clusters given a parameter for the minimum cluster size (in number of elements per cluster). This parameter indirectly affects the number of clusters. In order to determine the optimal minimum cluster size parameter for the OPTICS algorithm, we tested a range of minimum cluster sizes and computed an indicator of clustering quality (the mean silhouette score, see Methods). The highest silhouette score corresponded to a minimum cluster size of 8 in rodents as shown in 2B). With this choice of parameter, OPTICS identified five clusters (abbreviated as ‘R0-R5’), corresponding to five sub-types of synaptic dynamics.

The main characteristics of these functional subtypes can be seen by inspecting the shape of the SRP kernel, its baseline, and the parameter regulating the standard deviation (SD) of responses. Mean SRP kernels of each rodent cluster (Fig. 2C) and SD (Fig. 2D) revealed qualitatively distinct dynamics. Clusters ‘R3’ and ‘R4’ were characterized by kernels with large positive amplitude, which implements facilitation. These two clusters can be distinguished by considering the timescale of facilitation, which is faster for cluster ‘R4’. Clusters ‘R1’ and ‘R2’ are characterized by low baselines combined with kernels that have negative amplitudes and are associated with depressing synapses. These two clusters can be distinguished by the SD parameter, as cluster ‘R2’ was associated with the smallest degree of variability. Cluster ‘R0’ displayed a median kernel that is bimodal, which produces a slow and gradual form of facilitation, and can even produce depression for very fast stimulation.

**Fig. 2.**
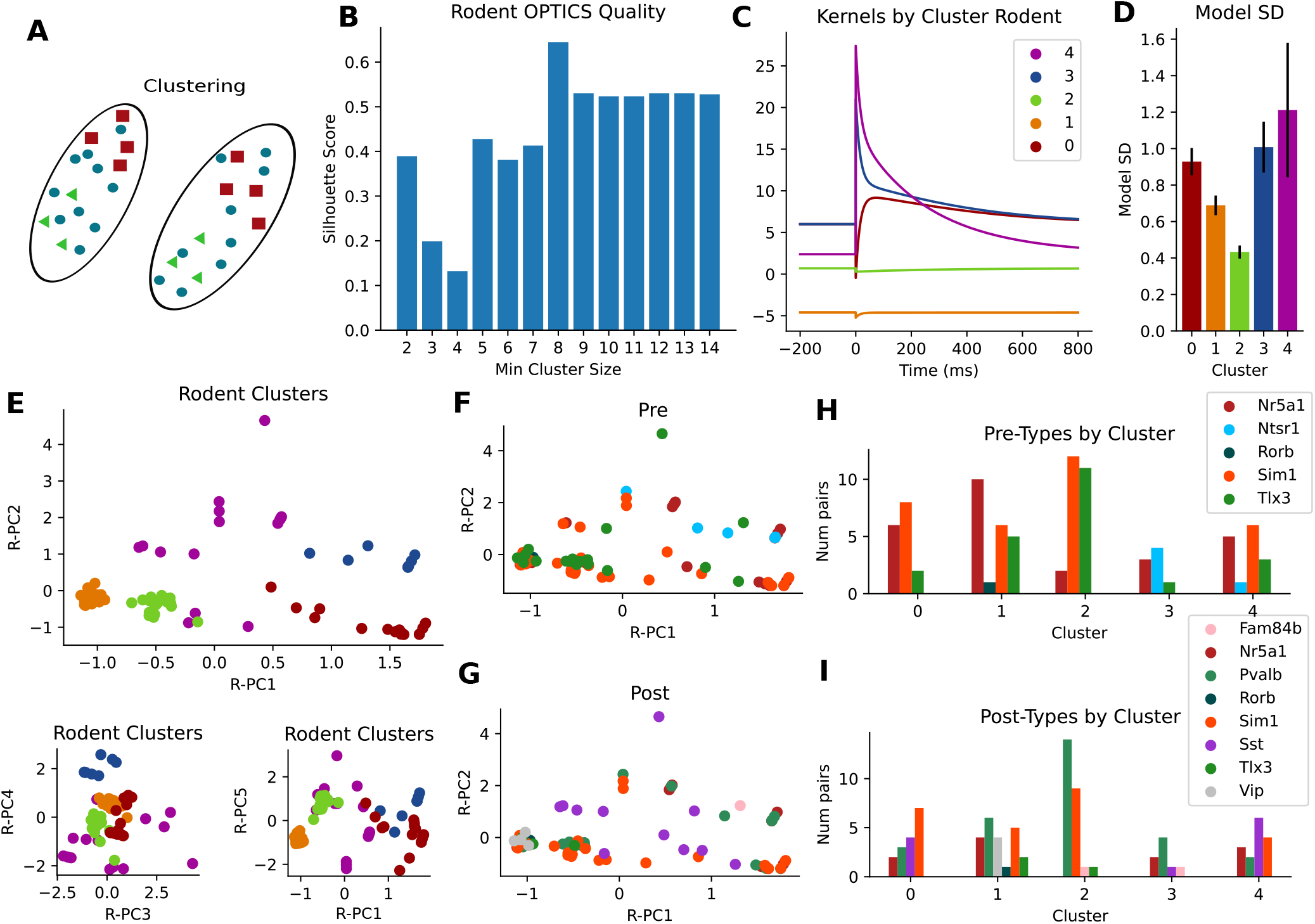
Unsupervised clustering reveals 5 clusters in rodent data. **A**: visual schematics of the process of unsupervised clustering. Synapses are grouped together based on model parameter similarity without taking labels such as cell subclass into account. **B**: silhouette coefficient, a measure of clustering quality, is shown against minimum cluster size. The maximum Silhoutte coefficient value corresponds to a minimum cluster size of 8 which was used for all rodent clustering. **C**: median kernels of each cluster are shown for each cluster based on the clustering of 86 glutamatergic synapses. **D**: Using the same color code as in C, model SD is shown for each cluster. Error bars show standard error of the mean (SEM). **E**: clusters generated by the OPTICS algorithm are plotted in different principal components of rodent data (R-PCs, where R-PC1 is the first principal component) in colours corresponding to B. **F-G**: In the same low-dimensional space as E (top), each dot represents a synapse coloured according to pre-synaptic subclass (F) or post-synaptic subclass (G). **H-I**: the number of synapses of each pre-synaptic subclass (H) and each post-synaptic subclass (I) is shown for each cluster with colours matched to C-E.

To visualize how these clusters relate with transcriptomic subclasses, we first plotted the model parameters for each pair projected into the space of their principal components (2E). As expected, each cluster occupied a restricted subregion of the space defined by the first two principal components. The median kernels and SD parameters of each colour coded cluster are shown in Fig. 2C-D. In the same space, we then labeled the pairs according to the pre-synaptic (Fig. 2F) or the post-synaptic (Fig. 2G) cell subclass. All clusters contained a mixture of cell subclasses both pre- and post-synaptically. This is consistent with the large heterogeneity of synaptic dynamics associated with transcriptomic subclasses (40, 47) (Fig. S1). Counting the number of pre-synaptic (Fig. 2H) or post-synaptic (Fig. 2I) subclasses found within each cluster shows that cell types tend to belong to specific functional subtypes. For instance, post-synaptic Pvalb tends to belong to cluster ‘R2’ (14 of the 29 connections) and cluster ‘R1’ (6 of the 29 connections), the two depressing subtypes. As another example, post-synaptic Sst tends to belong to cluster ‘R0’ (4 of the 11 connections) and cluster ‘R4’ (6 of the 11 connections), two facilitating subtypes. These observations are therefore consistent with the previous experimental reports of facilitating connections onto Sst and depressing connections on Pvalb-positive cells (16–18). Together, our results support a view where STP subtypes distribute in all cellular subclasses, but in subclass-specific proportions (46).

### Conserved functional subtypes are found in human synapses

A question of salience to this work is whether clusters emerging from our unsupervised process are consistent between species (Fig. 3A). We repeated the fitting and clustering procedure on human data from the Allen Institute Synaptic Physiology Dataset. The minimum cluster size was again selected to match the highest Silhouette Coefficient (minimum cluster size of 10, Fig. 3B). With this clustering parameter, OPTICS found 6 clusters. The similarity between clusters is illustrated in Figure 3C where the clusters of both species are plotted onto a set of principal components defined on the joint cross-species dataset (J-PC). The primary areas occupied by individual clusters are highlighted with clouds that are colour matched to closely corresponding clusters from the other species. Median kernels of the most similar clusters are shown next to each other, suggesting similar subtypes in the different species.

**Fig. 3.**
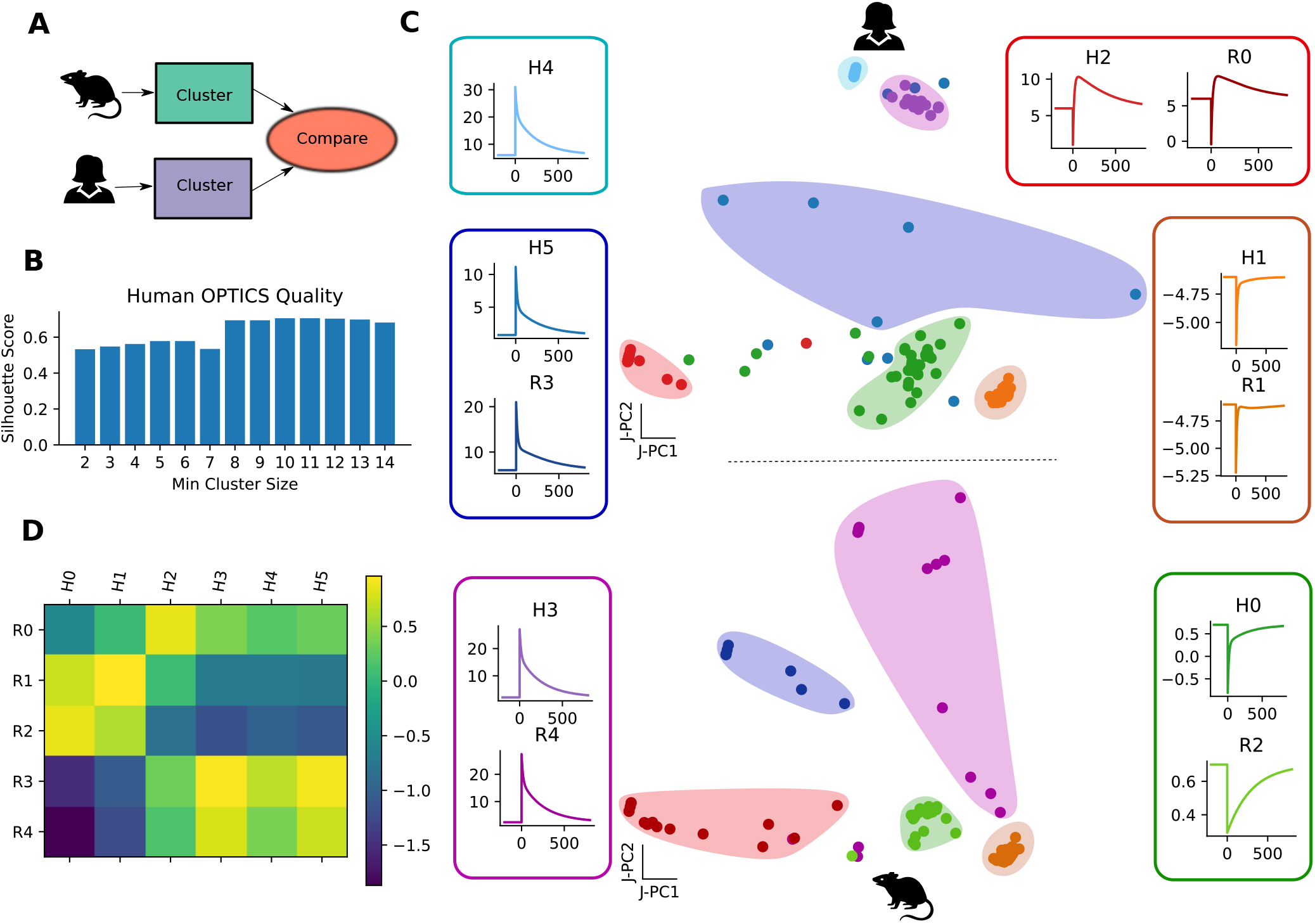
Unsupervised clustering shows functional alignment between rodent and human data. **A**: Visual schematic demonstrating that the same clustering protocol is applied to both human and rodent synapses and groups of functionally similar synapses are compared without using references to pre-defined types. **B**: As in Fig. 2B, different minimum cluster sizes are compared for human data based on Silhouette coefficient. The peak Silhoutte coefficient value corresponds to a minimum cluster size of 10 which was used for all human clustering. **C**: Scatter plot showing human (139 pairs) and rodent (86 pairs) clusters in the first jointly defined two principal component dimensions (J-PCs). The closest matches between human (ex: ‘H2’) and rodent (ex: ‘R0’) clusters are indicated with similarly coloured clouds and show strong qualitative alignment across species for all but one cluster. Corresponding median kernels are also presented. **D**: the similarity measure between the parameters of rodent and human clusters is shown as heat map (see Methods).

While one more cluster was found in the human synapses, this manifests itself through two clusters with very similar facilitating dynamics (‘H4’ and ‘H5’ in Fig. 3C). While relatively small, other notable differences exist, including stronger, but shorter lived depression in human cluster ‘H0’ than in its corresponding rodent cluster (‘R2’). To quantify the level of consistency between clusters, we computed a similarity measure incorporating a weighted combination of the temporal correlation of median kernels of each cluster and the difference in the modelled variance of their constituent synapses (Fig. 3D). In sum, although the exact number of subtypes may not be five (see Discussion), human synapses display similar functional modes to those found in the rodent data.

### Supervised learning relies on similar functional sub-types

Having demonstrated that neither the pre- nor the post-synaptic cell subclass aligns perfectly with one particular functional subtype, we wondered whether features of synaptic dynamics could predict molecular subclass. Despite imperfect alignment, it is still possible that the preponderance of specific functional subtypes within specific cell subclasses delineates such genetically-defined categories. To test this, we used supervised learning to predict cell subclass labels using only synaptic dynamics data (Fig 4A). We compared various machine learning algorithms as shown in Figure S4 and selected a multi-layer perceptron (MLP) with three hidden layers because it displayed the highest overall performance. The MLP was trained to predict cell subclass either from a set of physiological features (e.g. paired-pulse ratio, release probability, etc. See Methods) or from the SRP model parameters. In both cases, when predicting post-synaptic subclass, cross-validated accuracy was better than a ‘baseline’ classifier (31.76%) which always predicted the most frequent label in the training set (Fig. 4B) but this difference only reached significance in the case of the model representation (model accuracy 45.14%, P<0.001, phys accuracy 33.71%, P=0.297, bootstrap test for difference to baseline prediction).When predicting pre-synaptic sub-class, we found no significant difference with baseline prediction (36.16%) either using model parameters (41.36%, P=0.136, bootstrap test for difference with baseline) or using physiological features (33.71%, accuracy below baseline, Fig. 4B). Directly comparing classification accuracy using model parameters against using physiological features did not show a significant difference either in pre-synaptic (P=0.133, bootstrap test) or post-synaptic (P=0.074) subclass prediction. In summary, synaptic dynamics support some reliable predictions with respect to post-synaptic but not pre-synaptic subclass.

**Fig. 4.**
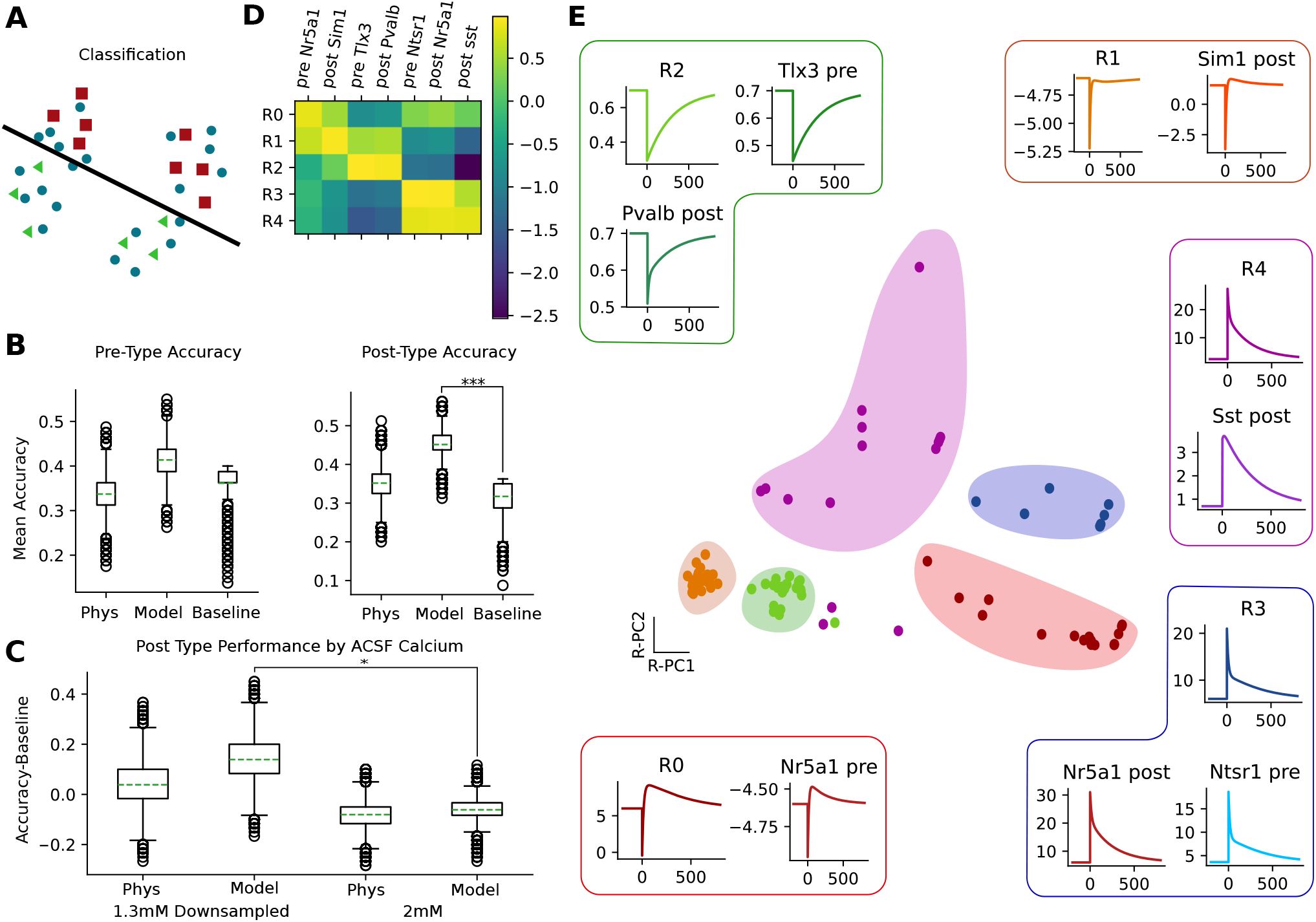
Clusters correspond to model features that are helpful for predicting pre- or post-synaptic cell identity. **A**: a visual schematic for illustrating that supervised learning separates data by optimally separating known subclasses, potentially at the expense of how the data is clustered. **B**: Mean cross-validation accuracy for predicting pre-type (left) and post-type (right) is shown for three models. The ‘baseline’ model uses the most frequently occurring label as the prediction. The ‘Phys’ uses a a multi-layer perceptron (MLP) with three hidden layers trained on physiological features. This classifier showed the best predictive performance in a classifier comparison shown in S4. The ‘Model’ used an MLP with the same configuration trained on model parameters. For pre-type prediction (left), the mean accuracy of the ‘Baseline’ is not outperformed by the ‘Model’ (P=0.136) nor ‘Phys’ (P=0.764). For post-type prediction (left), the mean accuracy of the ‘Baseline’ is outperformed by the ‘Model’ (P<0.001) but not ‘Phys’ (P=0.297). **C**: shows the difference in accuracy above baseline in post-synaptic subclass prediction based on data collected in 1.3 mM vs 2 mM. In this case the 1.3 mM dataset is downsampled to match the number of 2 mM exemplars remaining (67 after our data processing and quality controls). The ‘Model’ representation trained on 1.3 mM data outperformed an equivalent representation in 2 mM data (P=0.012) and the difference in performance between physiology representation in 1.3 mM and 2 mM trended towards significance (P=0.107). Histograms of differences resulting from the bootstrapping procedure used to generate these p-values are shown in S3. **D**: shows a similarity measure (see Methods) between unsupervised clusters and groups predicted by supervised learning incorporating temporal correlations between median kernels and the difference of mean variance parameters. These measures demonstrate alignment between functionally defined STP subtyes and the features relevant to the prediction of cell subclasses. Only the predicted subclasses with the highest measure of similarity to a rodent cluster are shown. **E**: Scatter plot of model parameters with colouring based on cluster identity (as in Fig. 2E). Insets highlight the qualitative similarity of median kernels of the different cell subclasses and functionally defined classes shown in D.

We next trained an MLP on physiological features and SRP model parameters of data obtained with a higher concentration of calcium in the bath (2 mM). In this case it was not possible for the MLP to outperform the baseline (Fig. 4C). However, there were more data available in physiological conditions (1.3 mM calcium) than in normal conditions (2 mM), so we controlled for the effect of dataset size by downsampling the amount of physiological calcium data to match that of the normal calcium. Even after downsampling it was still possible to perform above baseline in the prediction of post-synaptic subclass (P=0.040, difference from baseline, bootstrap test). We further controlled for the effect of different baseline prediction in normal and physiological conditions by directly computing the differences between model-based and baseline prediction accuracy in both conditions. We found that classifiers based on physiological calcium data outperformed those trained on normal calcium data when predicting post-synaptic subclass (P=0.012, 100,000 bootstrap shuffle for each, Fig. S2). In all cases the strongest predictive power was attained by combining our model-based representation with data collected in physiological calcium conditions.

We next compared the model features exploited by the MLP to the clusters obtained using the OPTICS algorithm. Specifically, the median kernel parameters and mean variance parameters of those synapses predicted by the MLP to belong to a particular subclass reveal features used by the machine learning algorithm to perform the classification. Similarity measures computed between such a sub-class stereotype and the representative parameters associated to a given cluster (‘R1’, ‘R2’, 舰) show selective alignment between cell subclass and clusters 4D. Each rodent cluster has strong correspondence with at least one specific pre- or post-synaptic subclass. In Figure 4E we illustrate the correspondence between label-aware and label-agnostic groupings by pairing the median kernels of the most similar groups from Figure 1D in distinct panels and highlighting the corresponding clusters in the space defined by the first two principal components of rodent SRP model fits. Together, our findings suggest that both label-aware and label-agnostic approaches find a similar set of relevant functional subtypes. Moreover, this alignment suggests that our clusters have particular relevance to cell subclasses.

## Discussion

This work uses machine learning techniques and mathematical modelling to infer distinct functional subtypes from the heterogeneity of cortical short-term plasticity in both humans and rodents. We found considerable overlap in inferred subtypes between human and rodent synapses and a strong alignment between our functional subtypes and several high level cell subclasses. Further, we have validated that these functional subtypes are not simply the product of noise by deliberately testing our classification in out of sample data. Our results also indicate that the dynamics of synapses belonging to different cell subclasses become less distinctive in calcium concentrations above physiological levels (ex: 2 mM).

Ultimately, are there really precisely 5 functional sub-classes of STP? Like all clustering approaches, our work is limited by the amount of data available, with more data typically enabling reliable identification of smaller clusters (44, 45, 50). What appears clear is that, consistent with previous work identifying multiple types of STD (37, 51, 52), the diversity in functional subtypes of STP is more nuanced than the simple division of STD and STF. Our results, particularly the convergence between human and mouse data, show that the basic dictionary of functional STP subtypes must contain at least five entries. Moreover, the partial overlap of these types with transgenically defined subclasses supports the hypothesis that STP properties cluster around genetically defined sub-types. Furthermore, our low-dimensional parameter space representation further supports a continuum of properties between some clusters (particularly the facilitating R0, R4 and R3, Fig. 2), but this variability can also arise from intrinsic experimental noise. Our data analysis can therefore not rule either in favor or against the presence of a continuum of STP properties between clusters.

What does the existence of our 5 to 6 subtypes change in our understanding of STP? A diversity of facilitating types suggests the possibility of multiple types of high rate coding schemes. Moreover, when these facilitating types are paired with the bimodal STP cluster we observed in both species, it hints at the possibility of decoding multilevel multiplexing schemes (53) wherein three or more different channels of information are encoded into a combination of long bursts, short bursts, and isolated events. Our STD subtypes were largely differentiated on the basis of variability which may indicate a difference in release probability. This may indicate a difference in the number of post-synaptic contacts which have been hypothesized to relate inversely to release probability as part of a mechanism to conserve energy at the synapse (54). We provide the full set of our fitted SRP model parameters labelled by group and by transgenically defined pre-synaptic and post-synaptic subclass in tables S3-S7 as a resource to aid the community in understanding the implications of these sub-types.

What determines if a connection belongs to a given functional subtype? We, like others (42), have observed that there is likely a transcriptomic basis for these sub-types, as clusters are aligned to features predictive for the subclass of either pre- or postsynaptic neuron. Given, however, the weak predictive power and the observation that each subclass has connections that belong to almost every cluster, we consider it likely that stronger correspondence between cell identities and functional dynamics will emerge from finer grain cell identity, such as at the type or sub-type level. If no finer grained transcriptomic basis is found, there remains the explanation that plasticity of STP properties is an important factor to explain their diversity (55).

These findings highlight several key takeaways for future experimental work. Since the most salient feature separating the STF subtypes is the timescale of facilitation, our results highlight the need for continued collection of cell-pair specific data containing stimulation protocols conductive to separating longer timescales. Relatedly, our supervised learning methods show better separation in near physiological bath calcium concentrations (4C), supporting its importance as an experimental parameter (13, 56, 57). In this case we theorize that higher bath calcium partially homogenizes STP, obfuscating meaningful cell identity specific dynamics. Lastly, our results support using a model-based representation instead of selected features of the recordings (Fig. 4B). Future experimental data can be integrated with our publicly available model parameters and model inference algorithm to help provide deeper insight about the dynamical nature of brain connectivity.

## Methods

### Spike Response Plasticity (SRP) Model

We modelled synaptic dynamics using the following flexible model from Rossbroich et al. (43) which is defined as follows:

Assume each pre-synaptic spike train gives rise to a post-synaptic current trace *I*(*t*) which corresponds to the sum of post-synaptic currents each triggered at a pre-synaptic time *t*_*j*_:

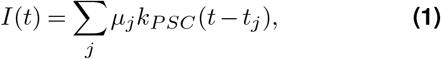

Where *μ*_*j*_ is the response to the jth spike in a normalized spike train and *k*_*PSC*_ is a kernel that describes a stereotyped PSC time course. We then defined an efficacy train *E*(*t*) as the multiplication of a time varying signaling *E*(*t*) which defines the synaptic efficacy at a given time with a spike train (*t*):

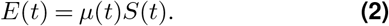

This allows us to rewrite the current trace as a convolution *I* = **k**_*PSC*_\**E* treating **k**_*PSC*_ as a kernel corresponding to the stereotyped PSC time course. We then model the efficacies *μ*_*j*_ as the output of a nonlinear readout of a linear filtering:

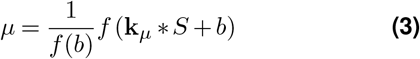

For this work, we select the kernel *k*_*μ*_ to be a sum of three exponential functions with time constants fixed to 15 ms, 200 ms, and 300 ms. The amplitudes of these functions and of the baseline parameter *b* are then free to be fitted for individual synapses providing a compact deterministic description of the dynamics of each synapse. We further describe the stochastic dynamics of the synapse by considering a sample of synaptic efficacies as a random variable *Y*_*j*_ with a mean value corresponding to the output of the deterministic description provided above:

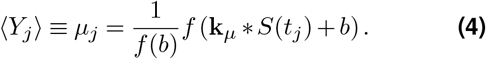

We then described the stochastic properties of the synapses by extracting the standard deviation *σ*_*j*_ from our mean squared error based fitting of mean synaptic behaviour. This allowed us to model the distribution of synaptic efficacies using a Gaussian distribution.

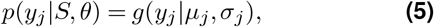

For a more extensive explanation and evaluation of this model see (43).

### Data Processing

We fit our data to a specific subset of the Allen Institute Synaptic physiology dataset (42) selected via the following criteria: Unless otherwise specified we selected data with a bath calcium concentration of 1.3 mM and bath magnesium concentration of 1 mM magnesium. The one exception is that for comparisons with 2 mM calcium data we selected data with 2 mM bath calcium. All queries required that the Allen Institute Synaptic Physiology dataset label a cell pair as a ‘synapse’, as excitatory, and not as a gap junction. Human or rodent data were specified for the respective analyses. We further excluded voltage clamp recordings, any pairs that were not associated with a ‘PulseResponseFit’, and any pairs with pre-synaptic labels corresponding to gabaergic interneuron types (Sst, Pvab, Vip). We then pooled run data with 1-3 ms variability in delay duration (125 ms, 126 ms, 127 ms, and 128 ms to 125 ms and 250 ms, 251 ms, 252 ms, and 253 ms to 250 ms) and padded runs missing responses to some stimuli with ‘nan’ values. In rodent data we excluded all synapses for which either presynaptic or postsynaptic cells were not transgenically identified. We also excluded synapses that lacked an experimental trail with 50 Hz induction frequency and 250 ms delay, which we used in calculation of our ‘physiological’ representation. We also excluded all synapses with paired pulse ratios over 4 to exclude a small number with unrealistically high values likely corresponding to an initial failure. On a protocol by protocol basis we excluded responses with initial response values above 0.01 volts and replaced any values below 10^−9^ volts with the value 10^−9^ volts to remove extreme values. We then normalized all values in a protocol by dividing them by the average of the first responses after the aforementioned exclusions and floor value replacements.

### Model Fitting

Once data had been pre-processed, we fit SRP model amplitudes, baselines, and standard deviations using simplical homology global optimization from the Scipy scientific computing library (58) version ‘1.6.2’. We selected amplitude and baseline parameters for each synapse by minimizing the mean squared error of our model prediction compared to recorded data across all runs and protocols. Finally, we computed the standard deviation for each synapse model based on the associated fitting errors. The baseline parameter was constrained to a value range between -4.5 and 6 while the value ranges of the amplitudes of three summed exponential functions of our kernel were constrained to -150 to 150, -1000 to 1001, and -3000 to 3000 for corresponding time constants of 15 ms, 200 ms, and 300 ms.

### Unsupervised Learning

Prior to clustering and supervised learning we applied PCA and scaling using the Scikit-learn package (59). Once the data were processed and scaled we ran the OPTICS (Ordering Points To Identify the Clustering Structure) algorithm from the Scikit-learn library (59) with a ‘sqeuclidean’ metric for minimum cluster sizes between 2 and 14. In both species we identified the minimum cluster size corresponding to the highest Silhoutte Coefficient (a measure of cluster quality). We then performed all further analyses using OPTICS with the minimum cluster size set to this default, the metric set to ‘sqeuclidean’ and all other parameters set to the defaults from Scikit-learn version ‘0.24.1’. In rodents the minimum cluster size we used was 8 while in humans we used a minimum cluster size of 10.

### Cluster and Group Similarity Measure

In order to calculate cluster similarity as in 3D and 4D we computed a combination of the correlation of median kernels and the difference in the Gaussian standard deviation parameter of the SRP model. Specifically we computed a Pearson correlation of median kernels and subtracted a value corresponding to the absolute difference of SD means between clusters divided by the standard deviation of the standard deviation SRP parameter across all fits. This resulted in a similarity metric where a value of 1 corresponds to perfect temporal correlation with zero difference in mean SD parameters and lower values correspond to increasingly dissimilar groups.

### Supervised Learning

In order to perform supervised learning on our data we generated two representations corresponding to our SRP model fits and a basket of traditional physiological measures. The SRP representation included the amplitudes of fitted kernels, the baseline parameter, and the standard deviation parameter for each synapse. The physiological representation included the mean summed value of the first eight stimuli, the mean ratio of the fifth response over the first response, the mean ratio of the second response over the first response (pair pulse ratio) for all runs with a 50 Hz induction frequency. Additionally, the physiological representation also included the release probability and the mean value of the last four responses in a train (those which follow the ‘delay’ divided by the mean of the first four for all protocols. We computed predictive accuracy in the prediction of all synapses onto each transgenically defined presynaptic or postsynaptic type that met the inclusion criteria described in in our data processing section. We also removed one synapse because it was the single remaining exemplar of its subclass and therefore could not be both trained and tested on.

#### MLP Parameters

We used the Scikit-learn package (59), version ‘0.24.1’ to implement a multilayer perceptron (MLP) classifier with three hidden layers of 50 units each and all other parameters set to the default configuration parameters of the package.

#### Comparison Classifier Parameters

We used the Scikit-learn package (59), version ‘0.24.1’ to implement gradient boosting, logistic regression, Adaboost, random forest, and a support vector machine classifiers for comparison with the selected MLP architecture. We set the maximum number of iterations for the logistic regression to 10000, and the number of estimators for gradient boosting to 500, otherwise all parameters were set to the default configuration parameters of the package.

#### Cross-Validation Protocol

In order to ensure that all synapses would be used in both model fitting (training) and for model assessment (testing) a standard cross-validation protocol was applied. The data corresponding to all excitatory rodent cells recorded in 1.3 mM calcium conditions were presented in one of two representations (SRP model parameters or traditional physiological measures), randomly shuffled, and then partitioned into subgroups of 10 samples each which dictated the number of folds. After partitioning, all but one fold was used to fit an untrained classifier and then the performance of that model was measured based on predictive accuracy on the remaining held out samples. This procedure was repeated for each fold to ensure testing on all available data. Further, the entire procedure was further repeated for 1000000 random shuffles. The random shuffling was applied to minimize order and grouping effects and repeated a large number of times to minimize variance in estimates of mean accuracy.

At each stage of the shuffling the difference between supervised predictive accuracy and accuracy baseline was computed and the distribution of these differences was used for significance calculations. We computed an accuracy baseline by using the ‘DummyClassifier’ class from Scikit-learn (59) to predict the most frequently occurring label from each training set for all entries in the corresponding test set and calculate the corresponding predictive accuracy. These accuracy baseline values provide a measure of how accurate a model could become by simply learning which subclasses occurred most regularly without learning anything about underlying synaptic dynamics. We computed the differences of mean cross validation accuracies between representations and against baseline for each shuffle. We then computed p-values by considering the proportion of shuffles in which representations outperformed baseline or each other.

## Acknowledgements

We thank Luke Campagnola for feed-back on the project as well as Zachary Friedenberger, Emerson Harkin, Leonard Maler, and Jérémie Lefebvre for helpful discussions. This work was supported by The Canadian Institutes of Health Research (CIHR):Project Grant RN38364 (RN and KT, providing salary to JB) and by the Natural Sciences and Engineering Research Council of Canada: Postgraduate Scholarships-Doctoral (JB, providing salary to JB). The funders had no role in study design, data collection and analysis, decision to publish, or preparation of the manuscript.

