## Supplemental for "Functional Subtypes of Synaptic Dynamics in Mouse and Human"

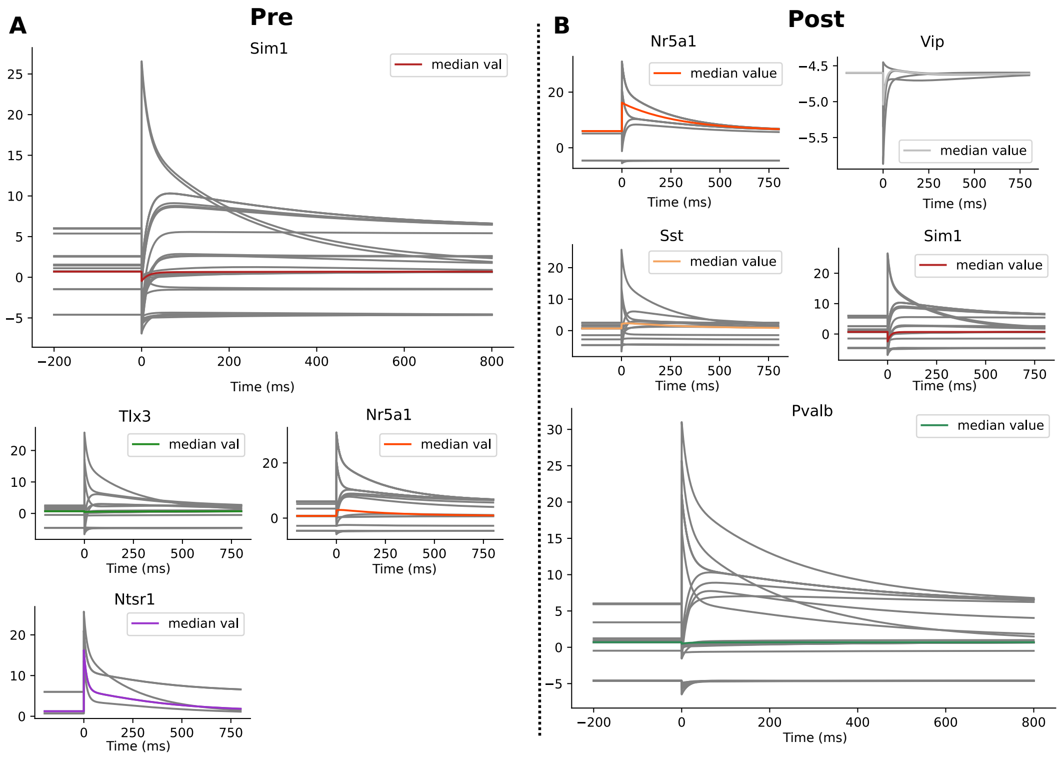


**Fig. S1.** Kernels of SRP model fits by pre-synaptic (**A**) and post-synaptic (**B**) cell subclass.

| Cluster # | Baseline | $A_1$ | $A_2$ | $A_3$ | SD |
|---|---|---|---|---|---|
| R0 | 5.996 | -150.0 | -1000.0 | 2570.635 | 0.929 |
| R1 | -4.6 | -9.419 | 46.448 | -67.945 | 0.689 |
| R2 | 0.7 | 0.0 | 0.5 | -122.902 | 0.432 |
| R3 | 6.0 | 150.0 | -1000.0 | 3000.0 | 1.008 |
| R4 | 2.39 | 150.0 | 1001.0 | 3000.0 | 1.211 |

**Table S1.** Median SRP parameters of rodent synapses used in clustering (Fig 2, Fig 3, and Fig 4 of 1.3mM Ca data labelled by pre-synaptic subclass, post-synaptic subclass, and cluster identity. The authors will provide fits to outstanding synapses from the Allen Institute Synpatic Physiology Dataset upon request.

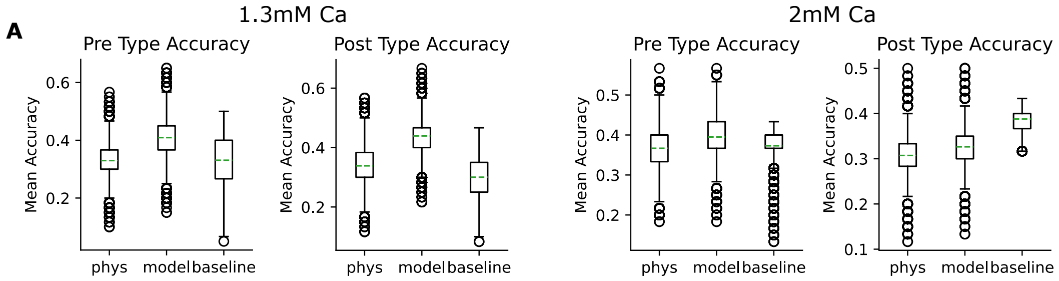


**Fig. S2.** Supervised accuracy by acsf calcium concentration matched for dataset size. When matched for size supervised accuracy is above baseline for model based pre- and post- type prediction in 1.3 mM Ca while neither representation supports accuracy above baseline in 2 mM Ca.

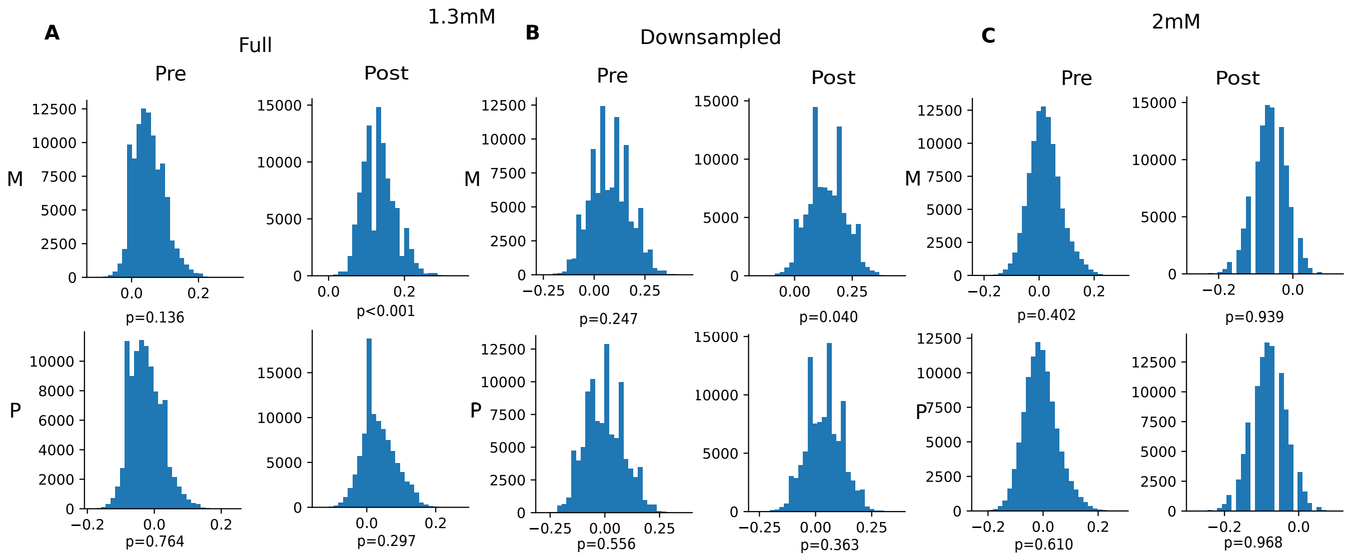


**Fig. S3.** Results by dataset and representation for full sized 1.3 mM ca (85 pairs), downsampled 1.3mM Ca (67 pairs), and full sized 2 mM Ca (67 pairs) of bootstrap testing for case when supervised model outperforms baseline.

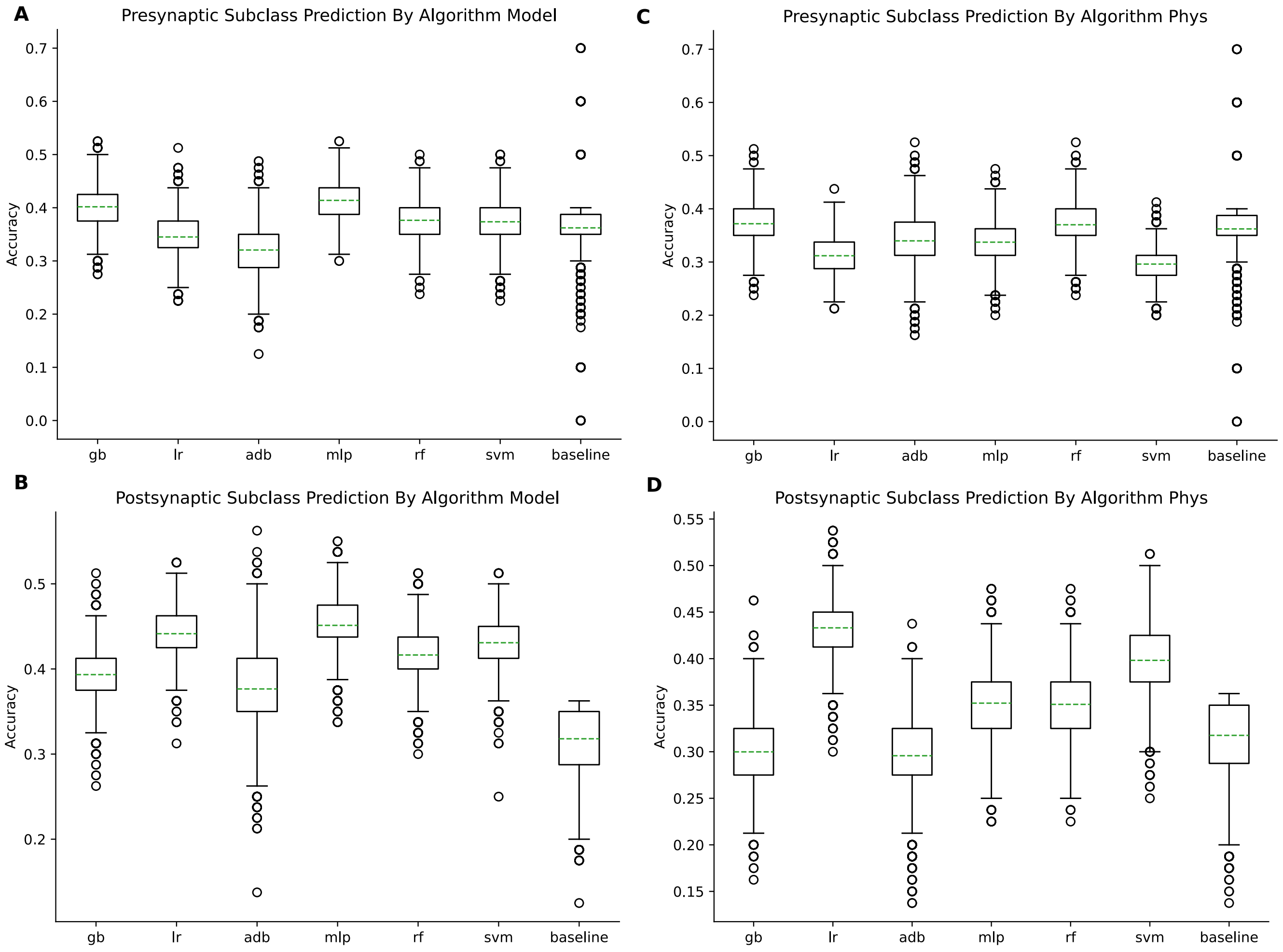


**Fig. S4.** Boxplots show supervised accuracy in prediction of transgenically defined pre-synaptic and postsynaptic subclass is shown for different algorithms across 10,000 bootstrap iterations. The green line represents the median accuracy while box lines denote upper and lower quartiles of the data. The algorithms are as follows: 'gb'=gradient boosting classifier, 'lr'=logistic regression, 'adb'=Adaboost classifier, 'mlp'=multi-layer perceptron, 'rf'=random forest, 'svm'=support vector machine. Finally 'baseline' displays the accuracy which can be attained by always predicting the most frequently occurring label from the training set. **A**: shows accuracies for prediction of pre-synaptic subclass using the model representation. **B**: shows accuracies for prediction of post-synaptic subclass using the model representation. **C**: shows accuracies for prediction of pre-synaptic subclass using a representation based on traditional measures. **C**: shows accuracies for prediction of post-synaptic subclass using a representation based on traditional measures.

| Cluster # | Baseline | $A_1$ | $A_2$ | $A_3$ | SD |
|---|---|---|---|---|---|
| H0 | 0.7 | -16.515 | 0.5 | -123.938 | 0.367 |
| H1 | -4.6 | -7.866 | -10.322 | -6.996 | 0.651 |
| H2 | 6.0 | -150.0 | -1000.0 | 3000.0 | 0.624 |
| H3 | 1.957 | 150.0 | 1001.0 | 3000.0 | 0.806 |
| H4 | 6.0 | 150.0 | 1001.0 | 3000.0 | 0.521 |
| H5 | 3.063 | 120.753 | 691.768 | 1387.867 | 0.758 |

**Table S2.** Median SRP parameters of Human synapses used in Fig 3 labelled by cluster identity.

| Cluster # | Pre-Subclass | Post-Subclass | Baseline | $A_1$ | $A_2$ | $A_3$ | SD |
|---|---|---|---|---|---|---|---|
| R0 | sim1 | sim1 | 1.559 | 150.0 | 1001.0 | 3000.0 | 0.684 |
| R0 | sim1 | sst | -1.453 | 25.938 | -4.456 | 82.471 | 2.635 |
| R0 | sim1 | sim1 | 1.11 | 150.0 | 1001.0 | 3000.0 | 1.229 |
| R0 | sim1 | sst | -4.6 | -7.624 | -99.418 | 230.763 | 3.346 |
| R0 | sim1 | sim1 | 5.378 | -150.0 | -78.406 | 186.272 | 0.817 |
| R0 | sim1 | sim1 | 2.615 | -150.0 | 131.654 | -59.945 | 0.854 |
| R0 | ntsr1 | pvalb | 0.7 | 150.0 | 1001.0 | 3000.0 | 1.725 |
| R0 | tlx3 | sst | 2.39 | -150.0 | 139.329 | 85.144 | 1.294 |
| R0 | nr5a1 | sst | -2.786 | -7.29 | 76.717 | 10.113 | 3.265 |
| R0 | tlx3 | sst | 2.543 | 150.0 | -76.716 | -38.671 | 1.211 |
| R0 | nr5a1 | nr5a1 | 6.0 | 150.0 | 1001.0 | 3000.0 | 0.759 |
| R0 | nr5a1 | nr5a1 | 6.0 | 150.0 | 1001.0 | 3000.0 | 0.572 |
| R0 | nr5a1 | nr5a1 | 6.0 | 150.0 | 1001.0 | 3000.0 | 0.911 |
| R0 | tlx3 | sst | 0.7 | 150.0 | 1001.0 | 3000.0 | 5.921 |
| R0 | nr5a1 | pvalb | 6.0 | 150.0 | 1001.0 | 3000.0 | 0.838 |
| R1 | sim1 | sim1 | 6.0 | -150.0 | -1000.0 | 3000.0 | 0.636 |
| R1 | sim1 | sim1 | 6.0 | -150.0 | -1000.0 | 3000.0 | 1.163 |
| R1 | sim1 | sim1 | 6.0 | -150.0 | -1000.0 | 3000.0 | 1.202 |
| R1 | sim1 | sim1 | 6.0 | -150.0 | -1000.0 | 2424.284 | 0.918 |
| R1 | nr5a1 | sst | 0.867 | -51.347 | -1000.0 | 1443.264 | 1.37 |
| R1 | sim1 | sim1 | 6.0 | -150.0 | -1000.0 | 2549.991 | 0.753 |
| R1 | nr5a1 | nr5a1 | 5.129 | -150.0 | -1000.0 | 2591.279 | 0.886 |
| R1 | sim1 | sim1 | 1.451 | -89.783 | -997.154 | 1742.645 | 1.203 |
| R1 | nr5a1 | pvalb | 5.992 | -150.0 | -1000.0 | 2466.657 | 0.79 |
| R1 | sim1 | sst | 0.7 | -7.317 | -775.689 | 1148.184 | 1.769 |
| R1 | nr5a1 | pvalb | 6.0 | -150.0 | -1000.0 | 3000.0 | 0.618 |
| R1 | nr5a1 | pvalb | 3.437 | -149.731 | -999.969 | 3000.0 | 0.891 |
| R1 | nr5a1 | nr5a1 | 6.0 | -150.0 | -1000.0 | 3000.0 | 0.939 |
| R1 | sim1 | sim1 | 6.0 | -146.122 | -1000.0 | 2362.117 | 1.002 |
| R1 | tlx3 | sst | 1.462 | -55.765 | -1000.0 | 1893.27 | 1.227 |
| R1 | tlx3 | sst | 1.761 | -125.759 | -997.179 | 2999.891 | 0.669 |
| R2 | sim1 | sim1 | -4.6 | -7.66 | -33.073 | 22.044 | 0.647 |
| R2 | tlx3 | vip | -4.6 | -3.32 | 110.956 | -180.818 | 0.69 |
| R2 | tlx3 | tlx3 | -4.6 | -9.227 | 70.218 | -77.672 | 1.011 |
| R2 | sim1 | sim1 | -4.6 | -9.966 | 61.042 | -93.411 | 0.804 |
| R2 | tlx3 | pvalb | -4.6 | -28.072 | 61.926 | -75.184 | 0.666 |
| R2 | sim1 | sim1 | -4.6 | -35.518 | 148.581 | -170.496 | 1.4 |
| R2 | tlx3 | tlx3 | -4.6 | -14.947 | -86.719 | 87.63 | 0.742 |
| R2 | sim1 | sim1 | -4.6 | 0.0 | 0.5 | -116.89 | 0.241 |
| R2 | tlx3 | vip | -4.6 | 0.945 | 59.943 | -63.886 | 1.291 |
| R2 | nr5a1 | pvalb | -4.6 | -13.503 | 83.353 | -129.803 | 0.688 |
| R2 | nr5a1 | vip | -4.6 | -17.28 | -48.275 | 37.563 | 0.631 |
| R2 | nr5a1 | vip | -4.6 | -12.531 | 73.997 | -84.775 | 0.893 |
| R2 | nr5a1 | pvalb | -4.6 | -8.847 | 16.271 | -37.019 | 0.676 |
| R2 | sim1 | sim1 | -4.6 | -14.363 | 74.233 | -118.549 | 0.554 |
| R2 | rorb | rorb | -4.6 | -5.261 | -40.707 | 43.983 | 0.859 |
| R2 | nr5a1 | pvalb | -4.6 | -2.579 | -14.769 | -10.98 | 0.809 |
| R2 | nr5a1 | nr5a1 | -4.6 | 1.984 | 14.588 | -48.356 | 0.654 |
| R2 | nr5a1 | nr5a1 | -4.6 | -9.611 | 51.307 | -71.329 | 0.75 |
| R2 | sim1 | pvalb | -4.6 | -16.601 | -0.183 | -34.109 | 0.487 |
| R2 | nr5a1 | nr5a1 | -4.6 | -13.93 | 55.726 | -98.678 | 0.367 |
| R2 | nr5a1 | nr5a1 | -4.6 | -0.403 | -41.43 | 25.49 | 0.658 |
| R2 | nr5a1 | pvalb | -4.6 | -6.081 | 41.588 | -64.56 | 0.695 |

**Table S3.** SRP parameters of rodent synapses used in clustering (Fig 2, Fig 3, and Fig 4 of 1.3mM Ca data labelled by pre-synaptic subclass, post-synaptic subclass, and cluster identity. The authors will provide fits to outstanding synapses from the Allen Institute Synpatic Physiology Dataset upon request.

| Cluster # | Pre-Subclass | Post-Subclass | Baseline | $A_1$ | $A_2$ | $A_3$ | SD |
|---|---|---|---|---|---|---|---|
| R3 | tlx3 | pvalb | 0.7 | 0.0 | 81.724 | -227.126 | 0.584 |
| R3 | tlx3 | pvalb | 0.996 | -16.631 | -38.685 | 0.0 | 0.481 |
| R3 | tlx3 | pvalb | 0.7 | 0.0 | 78.149 | -232.641 | 0.409 |
| R3 | tlx3 | pvalb | 0.7 | 0.0 | -61.59 | 0.0 | 0.596 |
| R3 | sim1 | sim1 | 0.7 | -55.882 | 41.074 | -177.421 | 0.209 |
| R3 | tlx3 | pvalb | 0.7 | 0.0 | 7.827 | -192.771 | 0.421 |
| R3 | tlx3 | pvalb | 0.7 | -5.863 | -92.541 | 19.642 | 0.473 |
| R3 | tlx3 | pvalb | 0.7 | 0.0 | 0.5 | -117.382 | 0.482 |
| R3 | tlx3 | tlx3 | 0.7 | 0.0 | 0.5 | -145.484 | 0.412 |
| R3 | tlx3 | pvalb | 0.7 | -33.845 | 0.5 | 0.0 | 0.949 |
| R3 | tlx3 | pvalb | -0.472 | -1.849 | 28.778 | -92.434 | 0.484 |
| R3 | nr5a1 | pvalb | 0.7 | 0.0 | 2.965 | -113.871 | 0.353 |
| R3 | sim1 | sim1 | 0.7 | -32.43 | 56.095 | -239.692 | 0.197 |
| R3 | nr5a1 | pvalb | 0.7 | -5.478 | -53.159 | 0.0 | 0.471 |
| R3 | sim1 | sim1 | 2.537 | -140.36 | 21.434 | 20.514 | 0.895 |
| R3 | sim1 | sim1 | 0.7 | -46.681 | 37.647 | -157.16 | 0.251 |
| R3 | sim1 | sim1 | 0.7 | 0.0 | -153.257 | -11.955 | 0.316 |
| R3 | sim1 | sim1 | 0.7 | 0.0 | 0.5 | -108.263 | 0.484 |
| R3 | sim1 | sim1 | -1.477 | -15.949 | 25.276 | -49.139 | 0.493 |
| R3 | sim1 | pvalb | 0.7 | 0.0 | 0.5 | -136.81 | 0.391 |
| R3 | sim1 | sim1 | 0.7 | 0.0 | 0.5 | -122.902 | 0.728 |
| R3 | tlx3 | fam84b | 0.7 | 0.0 | 0.5 | -138.224 | 0.26 |
| R3 | sim1 | pvalb | 0.7 | 0.0 | 36.796 | -207.478 | 0.373 |
| R3 | sim1 | pvalb | 0.7 | 0.0 | 0.5 | -139.493 | 0.432 |
| R3 | sim1 | sim1 | 0.7 | -75.063 | 134.539 | -218.485 | 0.422 |
| R4 | nr5a1 | nr5a1 | 6.0 | 150.0 | -1000.0 | 3000.0 | 1.288 |
| R4 | tlx3 | fam84b | 2.108 | 150.0 | -1000.0 | 3000.0 | 1.783 |
| R4 | ntsr1 | sst | 0.7 | 150.0 | -895.099 | 2206.568 | 1.523 |
| R4 | ntsr1 | pvalb | 6.0 | 150.0 | -1000.0 | 3000.0 | 0.712 |
| R4 | ntsr1 | pvalb | 1.236 | 150.0 | -1000.0 | 3000.0 | 1.041 |
| R4 | nr5a1 | pvalb | 6.0 | 150.0 | -1000.0 | 3000.0 | 0.975 |
| R4 | nr5a1 | nr5a1 | 6.0 | 150.0 | -1000.0 | 3000.0 | 0.648 |
| R4 | ntsr1 | pvalb | 6.0 | 150.0 | -1000.0 | 3000.0 | 0.66 |

**Table S4.** Table S3 continued.

| Cluster # | Baseline | $A_1$ | $A_2$ | $A_3$ | SD |
|---|---|---|---|---|---|
| H0 | 1.569 | -150.0 | 861.935 | -1095.588 | 0.284 |
| H0 | 4.565 | -150.0 | 541.701 | -393.597 | 1.429 |
| H0 | 0.7 | 150.0 | 1001.0 | 3000.0 | 2.866 |
| H0 | 5.289 | -150.0 | 841.835 | -824.329 | 0.686 |
| H0 | 6.0 | 150.0 | -1000.0 | 3000.0 | 0.563 |
| H0 | 5.21 | 0.0 | 77.598 | -565.331 | 0.548 |
| H0 | 0.746 | 150.0 | -1000.0 | 3000.0 | 3.265 |
| H0 | 3.806 | 150.0 | 1001.0 | 3000.0 | 0.433 |
| H0 | -2.486 | 91.506 | -1000.0 | 1600.999 | 8.204 |
| H0 | 2.32 | 150.0 | 1001.0 | 3000.0 | 1.659 |
| H0 | 4.027 | 0.0 | -999.432 | 1174.736 | 0.829 |
| H0 | 0.664 | 150.0 | 1001.0 | -2998.096 | 0.377 |
| H1 | 1.57 | 13.168 | 0.5 | -192.997 | 0.757 |
| H1 | 2.483 | -28.825 | -180.67 | 153.011 | 0.45 |
| H1 | 0.7 | -51.859 | 0.5 | -105.349 | 0.254 |
| H1 | 0.7 | 0.0 | 0.5 | -276.722 | 0.361 |
| H1 | 0.7 | 0.0 | 117.739 | -275.422 | 0.794 |
| H1 | 2.713 | 0.0 | 562.402 | -943.423 | 0.482 |
| H1 | 4.356 | -51.456 | -1000.0 | 1202.349 | 0.658 |
| H1 | 0.955 | -24.277 | 0.5 | 12.811 | 0.715 |
| H1 | 0.7 | -39.848 | 43.745 | -179.294 | 0.293 |
| H1 | 0.7 | -2.321 | -184.095 | 189.423 | 0.965 |
| H1 | 0.7 | 0.0 | 0.5 | -126.766 | 0.287 |
| H1 | 4.538 | -12.072 | -125.379 | -113.326 | 0.569 |
| H1 | 1.546 | -149.96 | 204.618 | -222.919 | 0.174 |
| H1 | 0.7 | -11.625 | -150.047 | 183.723 | 1.06 |
| H1 | 0.7 | 0.0 | -386.531 | -0.0 | 0.237 |
| H1 | 0.7 | -31.221 | 0.5 | -178.025 | 0.254 |
| H1 | 1.346 | -66.451 | -0.562 | -1.26 | 0.577 |
| H1 | 0.7 | 0.0 | 0.5 | -193.137 | 0.301 |
| H1 | 0.7 | 0.0 | 0.5 | -167.43 | 0.355 |
| H1 | 0.7 | -36.401 | 0.5 | -55.93 | 0.386 |
| H1 | 0.7 | 0.0 | 0.5 | -141.22 | 0.29 |
| H1 | 0.7 | 0.0 | 20.359 | -133.113 | 0.337 |
| H1 | -0.323 | -28.632 | 72.564 | -93.155 | 0.249 |
| H1 | 0.7 | 0.0 | 0.5 | -121.11 | 0.283 |
| H1 | 0.7 | 0.0 | 0.5 | -137.052 | 0.478 |
| H1 | 2.038 | -150.0 | 58.254 | -15.985 | 0.693 |
| H1 | 6.0 | -82.671 | -999.673 | 1804.921 | 0.463 |
| H1 | 0.7 | -31.295 | -52.6 | 4.062 | 0.332 |
| H1 | 0.7 | -84.998 | 197.325 | -277.285 | 0.234 |
| H1 | 0.7 | 0.0 | 0.5 | -198.24 | 0.42 |
| H1 | 0.7 | -21.604 | 0.5 | -116.349 | 0.373 |
| H1 | 0.7 | -104.894 | 201.939 | -275.962 | 0.261 |
| H1 | 0.7 | 0.0 | 0.5 | -147.234 | 0.442 |
| H1 | 0.7 | -59.266 | 18.042 | -129.225 | 0.242 |
| H1 | 0.7 | 0.0 | 0.5 | -138.277 | 0.211 |
| H1 | 0.7 | 0.0 | 0.5 | -136.415 | 0.672 |
| H1 | 0.7 | -15.986 | -81.601 | 0.0 | 0.498 |
| H1 | 3.933 | -26.556 | -999.333 | 1317.705 | 0.956 |
| H1 | 0.7 | -17.045 | -71.659 | 0.0 | 0.274 |

**Table S5.** SRP parameters of Human synapses used in fig 3 labelled by cluster identity.

| Cluster # | Baseline | $A_1$ | $A_2$ | $A_3$ | SD |
|---|---|---|---|---|---|
| H1 | 0.7 | -17.642 | 0.5 | -118.894 | 0.206 |
| H2 | -4.6 | -5.566 | -38.06 | 7.54 | 0.283 |
| H2 | -4.6 | -28.549 | 52.86 | -58.612 | 0.706 |
| H2 | -4.6 | -22.91 | 89.488 | -123.189 | 0.94 |
| H2 | -4.6 | -6.278 | -128.35 | 131.387 | 0.683 |
| H2 | -4.6 | -21.032 | -13.423 | 30.092 | 1.074 |
| H2 | -4.6 | -33.737 | 80.916 | -58.656 | 1.853 |
| H2 | -4.6 | 8.027 | -2.929 | -4.935 | 1.117 |
| H2 | -4.6 | -13.89 | -64.613 | 58.268 | 0.393 |
| H2 | -4.6 | -9.453 | -7.221 | -9.057 | 0.483 |
| H2 | -4.6 | -0.409 | 36.581 | -95.896 | 0.785 |
| H2 | -4.6 | -2.354 | -36.32 | -4.175 | 0.983 |
| H2 | -4.6 | -14.043 | -37.834 | 10.008 | 0.404 |
| H2 | -4.6 | 0.0 | 0.5 | -117.17 | 0.389 |
| H2 | -4.6 | -14.47 | -19.724 | 1.992 | 0.336 |
| H2 | -4.6 | -5.607 | 50.018 | -81.542 | 0.619 |
| H2 | -4.6 | -5.587 | -13.71 | -12.126 | 0.496 |
| H3 | 6.0 | -150.0 | -1000.0 | 3000.0 | 0.642 |
| H3 | 6.0 | -150.0 | -1000.0 | 3000.0 | 0.557 |
| H3 | 6.0 | -150.0 | -1000.0 | 3000.0 | 0.515 |
| H3 | 6.0 | -150.0 | -1000.0 | 3000.0 | 0.901 |
| H3 | 6.0 | -150.0 | -1000.0 | 3000.0 | 0.536 |
| H3 | 6.0 | -150.0 | -1000.0 | 3000.0 | 0.888 |
| H3 | 6.0 | -150.0 | -1000.0 | 3000.0 | 0.485 |
| H3 | 6.0 | -150.0 | -1000.0 | 3000.0 | 0.559 |
| H3 | 6.0 | -150.0 | -1000.0 | 3000.0 | 0.777 |
| H3 | 6.0 | -150.0 | -1000.0 | 3000.0 | 0.744 |
| H3 | 6.0 | -150.0 | -1000.0 | 3000.0 | 0.88 |
| H3 | 6.0 | -150.0 | -1000.0 | 3000.0 | 0.697 |
| H3 | 6.0 | -150.0 | -1000.0 | 3000.0 | 0.411 |
| H3 | 6.0 | -150.0 | -1000.0 | 3000.0 | 0.54 |
| H3 | 2.585 | -150.0 | -1000.0 | 3000.0 | 0.762 |
| H3 | 6.0 | -150.0 | -1000.0 | 3000.0 | 0.54 |
| H3 | 6.0 | -150.0 | -1000.0 | 3000.0 | 0.456 |
| H3 | 6.0 | -150.0 | -1000.0 | 3000.0 | 1.239 |
| H3 | 6.0 | -150.0 | -1000.0 | 3000.0 | 0.45 |
| H3 | 6.0 | -150.0 | -1000.0 | 3000.0 | 0.748 |
| H3 | 5.335 | -150.0 | -1000.0 | 2999.994 | 0.847 |
| H3 | 3.476 | -150.0 | -1000.0 | 2999.999 | 0.624 |
| H3 | 6.0 | -150.0 | -1000.0 | 3000.0 | 0.514 |
| H3 | 6.0 | -150.0 | -1000.0 | 3000.0 | 0.67 |
| H3 | 6.0 | -150.0 | -1000.0 | 3000.0 | 0.431 |
| H3 | 6.0 | -150.0 | -1000.0 | 3000.0 | 0.506 |
| H3 | 6.0 | -150.0 | -1000.0 | 3000.0 | 1.126 |
| H3 | 6.0 | -150.0 | -1000.0 | 3000.0 | 0.628 |
| H3 | 6.0 | -150.0 | -1000.0 | 3000.0 | 0.654 |
| H3 | 6.0 | -150.0 | -1000.0 | 3000.0 | 0.433 |
| H3 | 6.0 | -150.0 | -1000.0 | 3000.0 | 0.458 |
| H3 | 0.7 | 0.0 | -1000.0 | 1657.434 | 1.827 |
| H3 | 6.0 | -150.0 | -1000.0 | 3000.0 | 0.6 |

**Table S6.** Table S5 continued.

| Cluster # | Baseline | $A_1$ | $A_2$ | $A_3$ | SD |
|---|---|---|---|---|---|
| H4 | 1.813 | 150.0 | 1001.0 | 3000.0 | 0.494 |
| H4 | 0.7 | 150.0 | 1001.0 | 3000.0 | 1.213 |
| H4 | 1.774 | 150.0 | 1001.0 | 3000.0 | 1.075 |
| H4 | 0.7 | 150.0 | 1001.0 | 3000.0 | 1.273 |
| H4 | 2.131 | 150.0 | 1001.0 | 3000.0 | 0.482 |
| H4 | 2.683 | 150.0 | 1001.0 | 3000.0 | 0.382 |
| H4 | 1.466 | 150.0 | 1001.0 | 3000.0 | 0.977 |
| H4 | 1.841 | 150.0 | 1001.0 | 3000.0 | 0.997 |
| H4 | 2.921 | 150.0 | 1001.0 | 3000.0 | 1.503 |
| H4 | 1.037 | 150.0 | 1001.0 | 3000.0 | 0.516 |
| H4 | 2.581 | 150.0 | 1001.0 | 3000.0 | 0.749 |
| H4 | 3.508 | 150.0 | 1001.0 | 3000.0 | 0.393 |
| H4 | 2.072 | 150.0 | 1001.0 | 3000.0 | 0.625 |
| H4 | 2.215 | 150.0 | 1001.0 | 3000.0 | 0.863 |
| H5 | 6.0 | 150.0 | 1001.0 | 3000.0 | 1.005 |
| H5 | 6.0 | 150.0 | 1001.0 | 3000.0 | 0.403 |
| H5 | 6.0 | 150.0 | 1001.0 | 3000.0 | 0.546 |
| H5 | 6.0 | 150.0 | 1001.0 | 3000.0 | 0.889 |
| H5 | 6.0 | 150.0 | 1001.0 | 3000.0 | 0.729 |
| H5 | 6.0 | 150.0 | 1001.0 | 3000.0 | 0.505 |
| H5 | 6.0 | 150.0 | 1001.0 | 3000.0 | 0.401 |
| H5 | 6.0 | 150.0 | 1001.0 | 3000.0 | 0.487 |
| H5 | 6.0 | 150.0 | 1001.0 | 3000.0 | 0.79 |
| H5 | 6.0 | 150.0 | 1001.0 | 3000.0 | 0.637 |
| H5 | 6.0 | 150.0 | 1001.0 | 3000.0 | 0.473 |
| H5 | 6.0 | 150.0 | 1001.0 | 3000.0 | 0.606 |
| H5 | 6.0 | 150.0 | 1001.0 | 3000.0 | 0.536 |
| H5 | 6.0 | 150.0 | 1001.0 | 3000.0 | 0.583 |
| H5 | 6.0 | 150.0 | 1001.0 | 3000.0 | 0.501 |
| H5 | 6.0 | 150.0 | 1001.0 | 3000.0 | 0.534 |
| H5 | 6.0 | 150.0 | 1001.0 | 3000.0 | 0.401 |
| H5 | 6.0 | 150.0 | 1001.0 | 3000.0 | 0.39 |
| H5 | 6.0 | 150.0 | 1001.0 | 3000.0 | 0.467 |
| H5 | 6.0 | 150.0 | 1001.0 | 3000.0 | 0.508 |
| H5 | 6.0 | 150.0 | 1001.0 | 3000.0 | 0.754 |
| H5 | 6.0 | 150.0 | 1001.0 | 3000.0 | 0.436 |
| H5 | 6.0 | 150.0 | 1001.0 | 3000.0 | 0.508 |
| H5 | 6.0 | 150.0 | 1001.0 | 3000.0 | 0.779 |

**Table S7.** Table S5 continued.
